# A genome-wide CRISPR Screening identifies targets that drive Tolerogenic Dendritic Cells

**DOI:** 10.64898/2026.03.23.713621

**Authors:** Xiaoguang Li, Liang Chen, Tiankai Han, Megha Suresh, Fedik Rahimov, Chao Gao, Jing Wang, Renze Ma, Joshua D Stender, Kanstantsin Katlinski

## Abstract

Tolerogenic dendritic cells (TolDCs) are essential for immune tolerance and offer promise for treating autoimmune diseases. Despite the clinical evidence of their therapeutic potential, the key molecular pathways guiding their differentiation and tolerogenic phenotype remain elusive due to complex interactions identified in functional assays. Here we investigated the molecular profiles and regulatory programs underlying the functional status of tolerogenic dendritic cell populations in response to known tolerizing agents. We identified CD86 as a consistent and robust marker downregulated in tolerogenic state. Using CD86 blocking antibodies or CRISPR-mediated gene inactivation we demonstrated that CD86 is functionally required for TolDC-mediated suppression of T cell proliferation and cytokine secretion, establishing CD86 as both a consensus phenotypic and functional screening marker and a mechanistic regulator of tolerance. Leveraging CD86 as a scalable readout, we performed a pooled genome-wide CRISPR-Cas9 knockout screen to identify regulators of TolDC function. This approach uncovered UBE2L6 as a novel modulator that promotes the tolerogenic phenotype and restricts TolDC-mediated T cell activation. Mechanistically, UBE2L6 deficiency leads to coordinated upregulation of ISG15 and USP18, indicating a possible ISGylation-dependent pathway regulating CD86 expression and tolerogenic function. Together, this study identifies pathways that can be targeted to promote immune tolerance in immune-mediated inflammatory diseases.

**Graphical Abstract:** 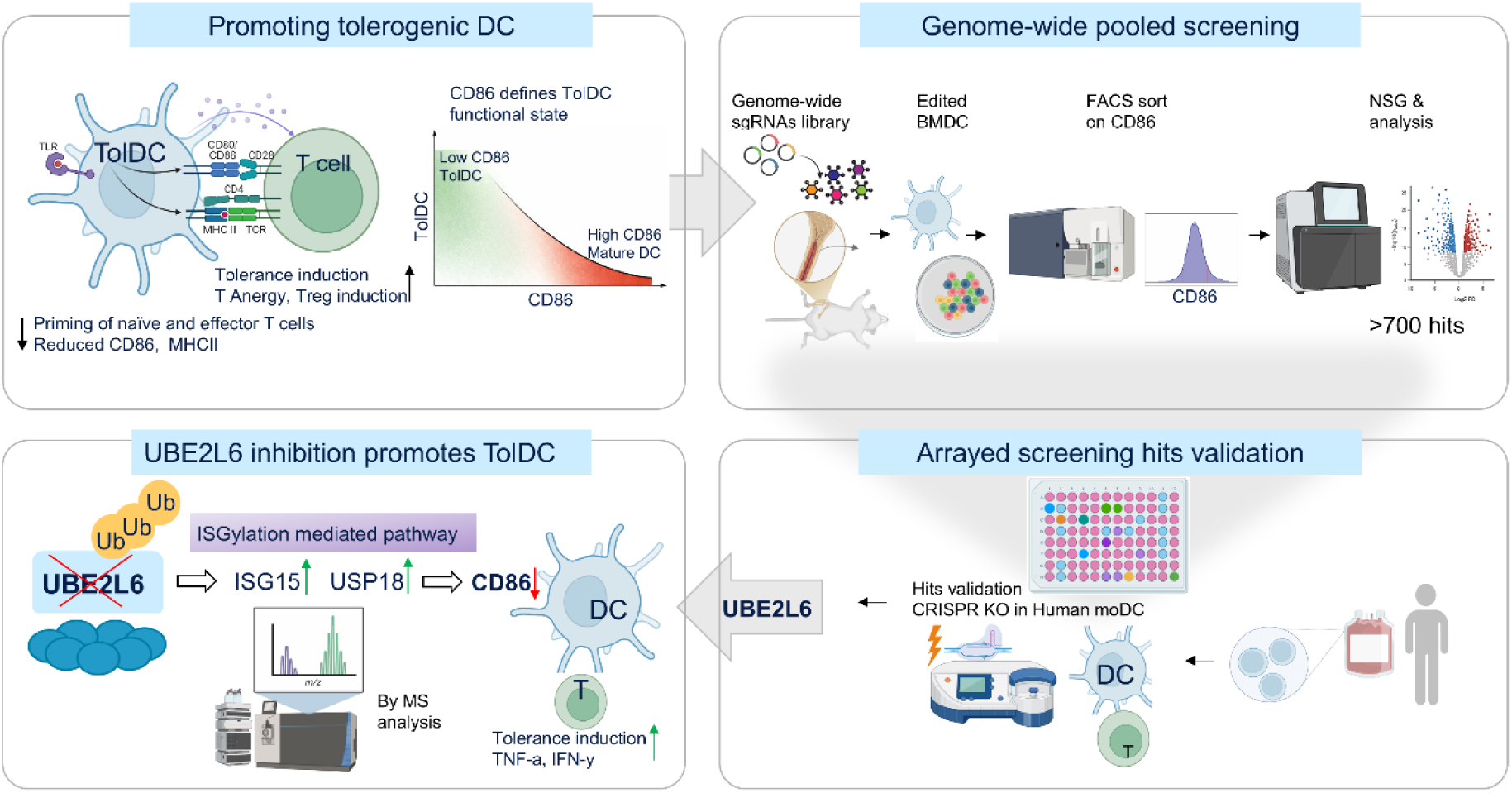

## Introduction

Autoimmune diseases arise from a breakdown of immunological tolerance mechanisms in which a reaction to self-antigens results in the generation of effector cells that can lead to tissue and organ damage^1–3^. Responses to “self” are typically a result of lymphocytes that have escaped central and peripheral tolerance that would normally prevent their affinity for self-antigens^4,5^. Thus, understanding mechanisms of tolerance and its role in immune dysfunction is a key to developing novel therapeutic strategies that aim to rebalance the immune system. The standard of treatment for autoimmune diseases has long been a broad spectrum of immunosuppressive agents and immunomodulatory biologics that are usually accompanied by long term side effects, life-long drug administration, and variable disease resolution^6–8^. Additionally, 30-50% of patients fail to respond to current therapies ^9–12^. Restoring immune tolerance therefore represents a transformative approach for the treatment of autoimmune diseases that can not only reduce the need for disease-modifying therapy but also offer prolonged improvement of clinical outcomes^13^.

Dendritic cells (DCs) are important regulators of immune homeostasis and activation, maintaining both central and peripheral tolerance and serving as a bridge between innate and adaptive immunity^14–16^. Tolerogenic dendritic cells (TolDCs) can inhibit memory and effector T cell responses due to their reduced expression of co-stimulatory markers, such as CD80, CD40, and CD86, as well as their enhanced ability to produce anti-inflammatory cytokines like IL-10. 0^17–19^. In general, these characteristics contribute to their ability to modulate immune responses, suppress effector T cell functions, and generate regulatory T cells (Tregs)^20^. As a result, reprogramming dendritic cells (DCs) into TolDCs has emerged as a promising cell-based therapy for the treatment of autoimmune diseases. In the clinic, TolDCs can be induced by immunosuppressive agents such as corticosteroids, dexamethasone, and vitamin D3; by anti-sense oligonucleotides targeting the CD40, CD80 and CD86 co-stimulatory molecules; or by Immunosuppressive cytokines or by genetic modifications ^21–25^. However, different TolDCs exhibit distinct features and properties and, current approaches to evaluate DC tolerogenicity rely on time-consuming and often unreliable methods such as phenotyping and allogeneic T cell proliferation assays^26^. These limitations highlight the need for appropriate selection of consensus TolDC biomarkers. While reduced expression of co-stimulatory molecules is linked to a tolerogenic state in dendritic cells, it is still unclear whether these molecules act solely as phenotypic markers of tolerance or play a direct, mechanistic role in the function of tolerogenic dendritic cells (TolDCs). Here, we generated functional TolDCs using multiple benchmark tolerogenic agents and confirmed reduced CD86 expression across TolDCs induced by different agents and its dependency in inducing tolerance.

Early-phase clinical studies have demonstrated feasibility and safety of ex vivo–generated TolDCs for autoimmune diseases such as type I diabetes and rheumatoid arthritis. In these studies, DC precursors are isolated from the patient, treated *ex vivo* to generate TolDC, loaded with autoantigens (optional), and administered back into the patient ^27^. Yet broader application of TolDC has been limited because the molecular mechanisms underlying tolerance remain elusive^28^. Current CRISPR technology provides opportunities for comprehensive analysis of biological pathways using pooled genome-wide libraries. However, these approaches require robust biomarkers that accurately reflect the biological process to serve as effective phenotypic readouts. Technical limitations in earlier screens often restricted analyses to subsets of prioritized genes^29^ or relied on immortalized cell lines^30^. Notably, most prior target identification efforts, including CRISPR-based screens in dendritic cells, have focused on uncovering targets that promote immunogenic functions in conventional DCs to help overcome immunotolerance^31,32^. We have developed an approach to identify novel targets that enhance the tolerogenic functions of dendritic cells by screening candidate genes in tolerogenic DCs using CD86 expression as a phenotypic readout. CD86 is a well-known co-stimulatory signal for T cell activation. reduced expression of CD86 was shown to be associated with tolerogenic DC cell phenotype and its silencing in DCs induced immune tolerance. Our screening recovered many of the key regulators involved in the TLR signaling and inflammatory responses, including NFKB1, TRAF6 and TIRAP. Through functional validation and quantitative proteomics, we identified a previously unrecognized UBE2L6 –ISG15–USP18 regulatory axis that controls CD86 expression and DC mediated T cell activation. Post-translational modification pathways, including ISGylation, are important modulators of innate immune signaling and antigen-presenting cell function. While components of the ISGylation machinery have been reported in inflammatory and antiviral responses, their role in DC tolerogenic programming has not been defined^33,34^. Altogether, our studies provide a comprehensive profile of key modulators that enhance TolDC function, deepening our understanding of strategies to establish TolDCs as a promising therapeutic approach for restoring immune tolerance.

## Results

### Tolerogenic DCs exhibit a reduced capacity to promote T cell proliferation and inflammatory cytokine production

We generated TolDCs using three well-characterized tolerogenic agents: vitamin D3 (VitD3), dexamethasone (Dex), and rapamycin (Rapa). The ability of TolDC to suppress T cell proliferation and inflammatory cytokines was evaluated in functional assays with human monocyte-derived DCs (moDCs) or murine bone marrow–derived DCs (BMDCs). Across all conditions, TolDCs displayed a reduced ability to induce T cell proliferation compared with control immunogenic DCs. CFSE dilution assays revealed a significant decrease in the frequency of proliferating CD4⁺ and CD8⁺ T cells following coculture with VitD3-, Dex-, or Rapa-induced TolDCs (Figure S1A-D). In addition, there was a induction of regulatory T cells (Figure S1E). Cytokine analysis of coculture supernatants demonstrated that TolDCs induced substantially lower levels of T cell cytokines, including IL-2, IFN-γ and TNF-α (Figure S1F). This impaired capacity was also observed in mouse systems in murine bone marrow–derived dendritic cells (BMDCs) (Figure S2). Together, these data confirm that TolDCs generated using distinct pharmacological agents are functionally characterized by attenuated T cell activation and inflammatory cytokine production in both human and murine assays.

### Consensus tolerogenic markers were identified across TolDCs induced by different agents

Although VitD3, Dex, and Rapa induce TolDCs through mechanistically distinct pathways, we hypothesized that these cells share a common transcriptional and phenotypic profile. To identify consensus tolerogenic markers, we performed comparative phenotypic profile across TolDCs generated with each agent. A panel of candidate markers—including CD40, CD86, CD80, IDO-1, MHCII or HLA-DR—were examined. Flow cytometric analysis confirmed that the costimulatory molecule CD86 was consistently and robustly downregulated across VitD3-, Dex-, and Rapa-induced TolDCs compared with control DCs, in both human monocyte-derived DCs and murine BMDCs (Figure 1). Taken together, CD86 is a robust and consensus biomarker that is highly associated with TolDC function.

**Figure 1:**
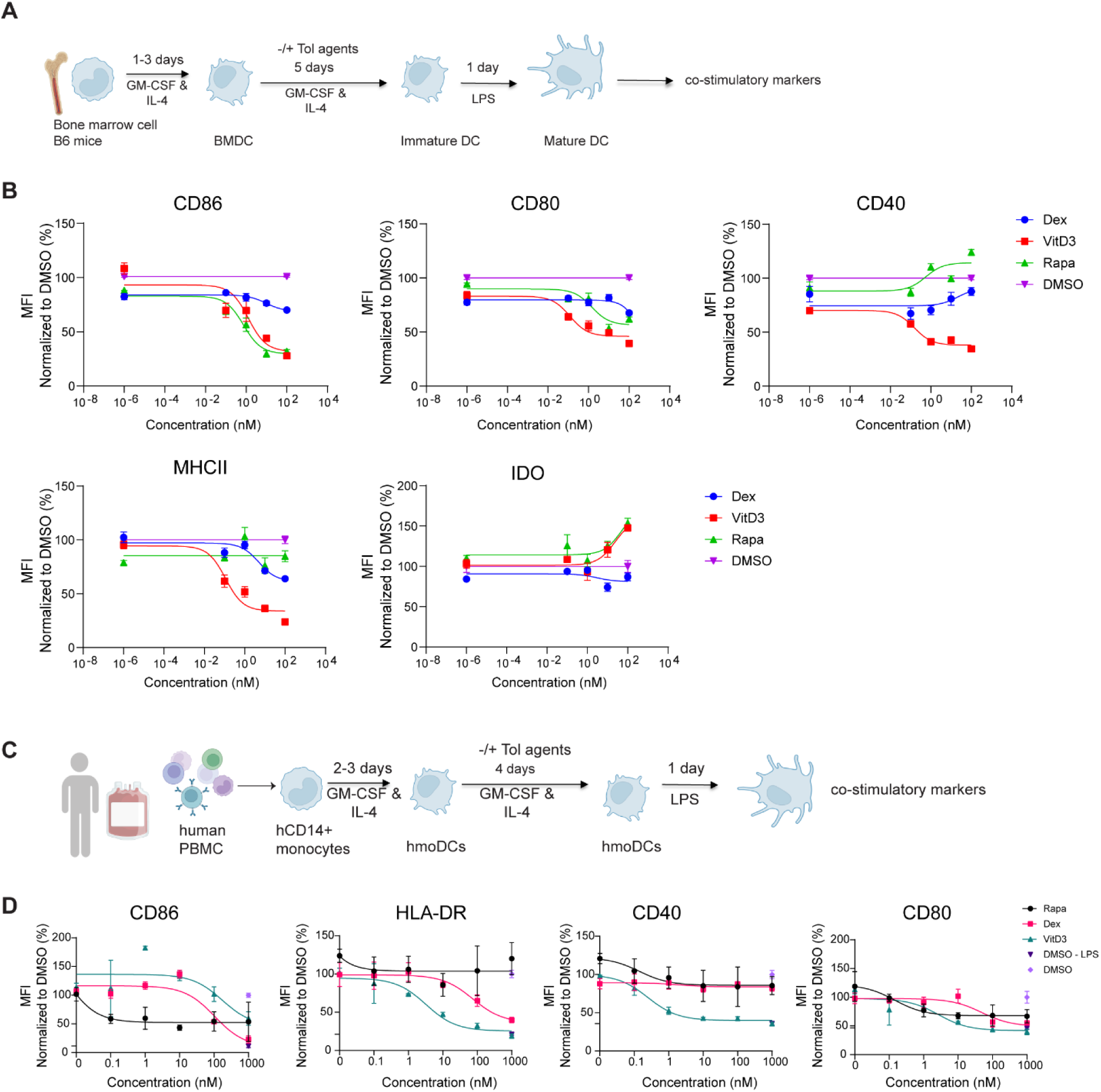
Identification of TolDCs consensus biomarkers in response to tolerizing agents Dex, VitD3 and Rapa. **(A)** Schematic representation of generation and characterization of mouse BMDCs and TolDCs. **(B)** Flow cytometry analysis of CD86, CD40, CD80, MHC-II and IDO expression in mouse tolDCs after treatment with indicated concentrations of tolerizing agents. **(C)**. Schematic representation of generation and characterization of human TolDCs. **(D)** Flow cytometry analysis of CD86, CD40, CD80, and HLA-DR in human tolDCs after treatment with indicated concentrations of tolerizing agents. The results represent N=3 donors or 3 independent experiments.

### Phenotypic marker CD86 reflects the functional tolerogenic capacity of TolDC and provides a robust phenotypic readout for genome wide screening

Next, to address whether CD86 contributes to TolDC function and its dependency in inducing tolerance, we knocked out CD86 using CRISPR RNP mediated KO in moDCs (Figure 2A) and characterized moDC differentiation overtime by assessing DC-SIGN and CD14 expression (Figure S3A). Analysis of cytokine profiles revealed increased anti-inflammatory IL-10 level in CD86-deficient moDCs, with no observed changes in pro-inflammatory cytokines TNF-α and IL-6 (Figure 2B). Next, co-culture of moDC with allogenic T cells demonstrated significantly reduced T cell proliferation and T cell cytokines, including IFN-γ, IL-2, IL-5, IL-13, and TNF-α in 4 out of 16 tested donor pairs (Figure 2C-F). Parallel studies in BMDC using CD86 blocking Abs consistently confirmed that induction of tolerance by TolDCs depends on CD86 inactivation, as evidenced by suppressed T cell proliferation and reduced production of inflammatory cytokines. (Figure 2G-I). Collectively, we identified CD86 as a consensus phenotypic and robust functional screening readout that contributes and reflects the tolerogenic potential of DCs.

**Figure 2:**
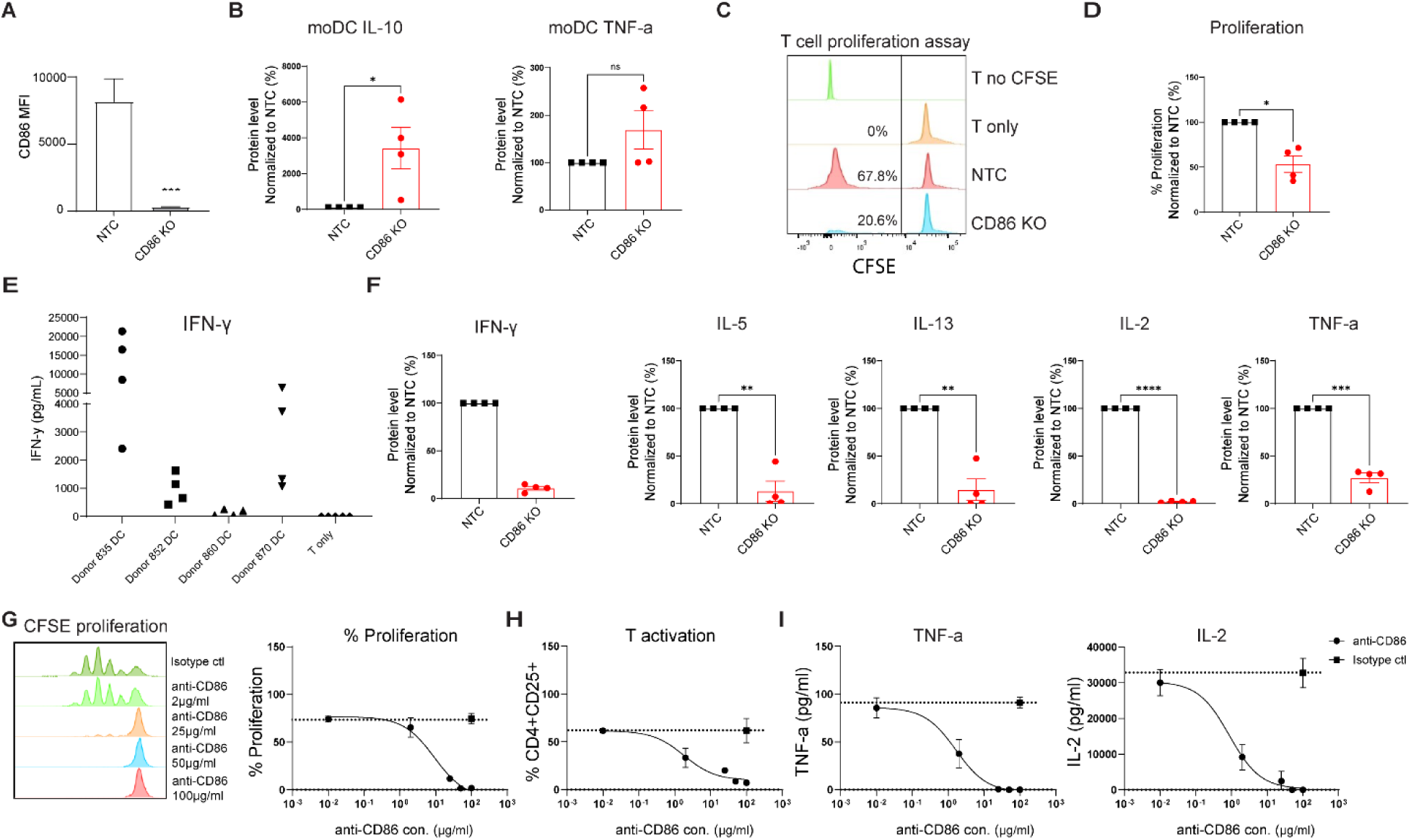
CD86 as a phenotypic biomarker linked to the functional status of TolDC. **(A)** CD86 was knocked out in moDCs using RNP/sgRNA complexes targeting CD86 (CD86 KO), with non-edited cells as a non-targeting control (NTC). Flow cytometer analysis of CD86 expression in NTC and CD86 KO moDCs. **(B)** Secreted cytokine levels of IL-10 and TNF- α by moDCs were measured by MSD and normalized to NTC (%). **(C–D)** T cell proliferation in the moDCs:T cell co-culture mixed lymphocyte reaction (MLR) assays. **(E)** Secreted IFN-γ level in the co-culture across 16 DC : T cell donor pairs, each dot representing one donor pair. **(F)** Secreted cytokine levels of IFN-γ, IL-5, IL-13, IL-2, TNF-α in co-culture supernatants were measured by MSD and normalized to NTC (%). **(G)** Flow cytometer analysis of T cell proliferation in CFSE assay. **(H)** Flow cytometer analysis of T cell activation was performed by measuring CD4+CD25+ cell frequency. **(I)** Secreted TNF-α, and IL-2 levels in the co-culture system treated with indicated concentrations of CD86 blocking antibody. CD86 expression, T cell activation, and proliferation (CFSE), assessed by flow cytometry; cytokines by MSD assay. Data from N=4 donors (each dot representing 1 donor or donor pair); statistical significance calculated by paired two-tailed t test; error bars indicate mean ± SEM; *p < 0.05; **p < 0.01; ***p < 0.001; ****p < 0.0001.

### Genome wide CRISPR-Cas9 screen in primary dendritic cells identifies novel targets that induce tolerogenic DCs

Next, we used CD86 as the screening readout for the genome-wide screen to identify TolDC regulators. We optimized the screen conditions in BMDCs by using a low multiplicity of infection (MOI = 0.1) and maximizing cell expansion to maintain library representation (Figure S3B).

Under these conditions, we performed pooled genome-wide CRISPR-Cas9 screen using a library of lentiviruses harboring 80,600 sgRNAs targeting 21,300 annotated protein-coding genes in a tolerogenic context. TolDCs were subsequently sorted based on high or low CD86 surface expressions, and sgRNAs enrichment and depletion in each population were measured by deep sequencing (Figure 3A-B, and Figure S3C-D). We achieved efficient gene knock down and identified 450 putative positive regulators and 345 negative regulators of the TolDC phenotype. Among the positive regulators were previously reported genes in the NF-κB, MAPK, or JAK-STAT signaling pathways, such as MAP3K14, NFKB1, TRAF6, and IFNAR1, and negative regulators included SOCS6 and PPM1D (Figure 3C-E). KEGG Pathway analysis further confirmed and revealed several well-established immune regulatory pathways, such as Toll-like receptor signaling pathway, IL-17 signaling pathway, MAPK signaling pathway, NF-κB signaling pathway, as well as metabolic pathways (Figure 3F). Gene Ontology enrichment analysis revealed the involvement of several immune-related pathways, including TNF receptor binding, MAPK activity, vitamin transmembrane transporter activation, and integrin binding. Additionally, we identified novel pathways such as ubiquitin binding and ubiquitin-like protein ligase activity (Figure S3E-F).

**Figure 3:**
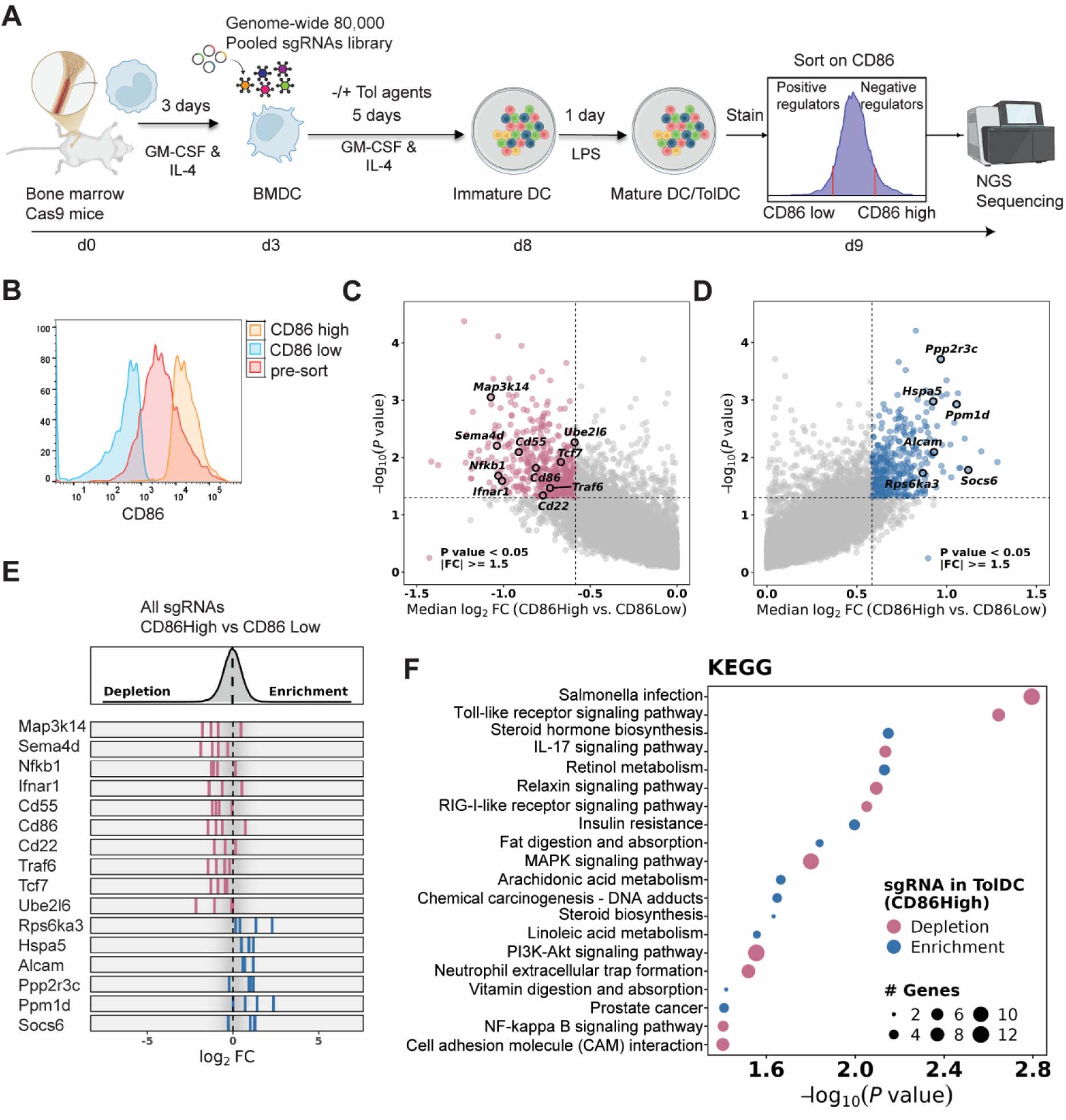
Genome wide CRISPR-Cas9 screen identifies novel targets that induce tolerogenic DCs. **(A)** Schematic representation of the genome-wide CRISPR-Cas9 screen in mouse BMDC. **(B)** Flow cytometer results for pre-sort, CD86 high and CD86 low samples prior to next generation sequencing. **(C–D)** A volcano plot of the hits from the screen. The x axis indicates the median log2 (fold change) (LFC) for all sgRNAs per gene, whereas the y axis depicts −log10 (P value). Statistically significant genes were identified by aggregating differentially ranked sgRNAs at the gene level for depletion or enrichment using the robust rank aggregation (RRA) method, together with a threshold of median LFC > 0.585. Positive regulators of CD86 are shown in red, and negative regulators are in blue. **(E)** Distribution of median LFC of sgRNAs for selected hits in CD86high versus CD86low cells. **(F)** Selected KEGG pathways enriched in DC86 positive (red) and negative (blue) regulators ranked by - log2(P value).

### Genetic associations of CRISPR screening hits with immune-mediated diseases

CRISPR screens identify genes that functionally regulate immune cell activation or tolerance, while pathway enrichment analysis groups these hits into biological pathways, aiming to identify the underlying mechanisms and prioritize key regulatory networks. Further integration of screening hits with genome-wide association studies (GWAS) of common immune-mediated diseases provides a unique approach for elucidating causal mechanisms and identifying candidate therapeutic targets^35^. Here, we leveraged the Open Targets Platform’s ‘locus-to-gene’ (L2G) model to systematically evaluate the genetic association of CRISPR screen hits that modulate CD86 expression and tolerogenic dendritic cell activity with human diseases. By integrating results from 129 case-control GWAS, alongside FinnGen and UK Biobank datasets covering 28 harmonized autoimmune and inflammatory phenotypes, we found that a significant subset of our screen genes were robustly associated with increased disease risk (Figure 4). Importantly, these associations spanned a variety of autoimmune disorders, suggesting that altered dendritic cell co-stimulatory signaling and tolerance pathways may underlie a common pathogenic mechanism of many immune-mediated diseases. Our findings directly link molecular regulators of the tolerogenic DC phenotype to genetic risk factors for immune-mediated conditions, providing new mechanistic insight and highlighting promising targets for future investigation.

**Figure 4:**
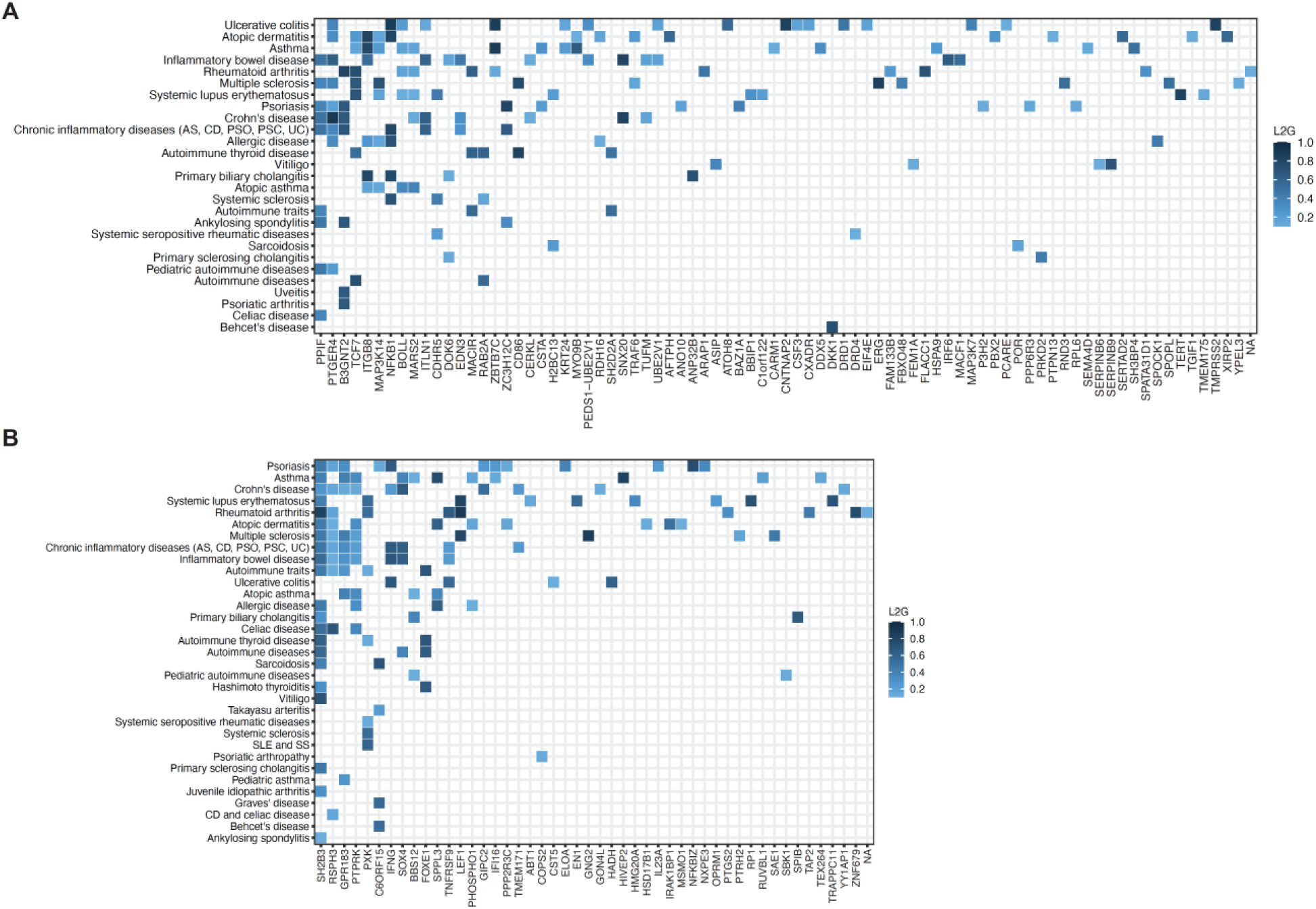
Genetic association of CRISPR screening hits with immune-mediated diseases. **(A)** Top locus-to-gene (L2G) scores from Open Targets Genetics for gene-disease pairs. Positive regulators identified in the CRISPR screen were linked to various immune-mediated diseases in GWAS with varying levels of confidence as determined by the L2G scores. **(B)** Negative regulators identified in the CRISPR screen were linked to various immune-mediated diseases in GWAS with varying levels of confidence as determined by the L2G scores. In both plots L2G scores above 0.1 are shown.

### Arrayed screen validated novel regulators driving tolerogenic DCs

To further confirm the screen findings and assess translational relevance to human systems, we validated selected hits in human monocyte-derived dendritic cells (moDCs). Given the therapeutic potential of enhancing immune tolerance in autoimmune diseases, we focused downstream analyses on the positive regulators. Knockdown of these candidates consistently significantly reduced CD86 expression (Figure 5A-B). Most hits also decreased HLA-DR expression, while having minimal effects on CD80 and CD83 (Figure 5C and Figure S4). In addition, these perturbations did not affect DC differentiation, as measured by DC-SIGN expression (Figure 5D). Together, these results indicate that the identified regulators specifically control CD86 expression without broadly impairing DC differentiation, supporting their potential role in selectively promoting tolerogenic DC programs and their applicability as therapeutic targets for autoimmune diseases.

**Figure 5.**
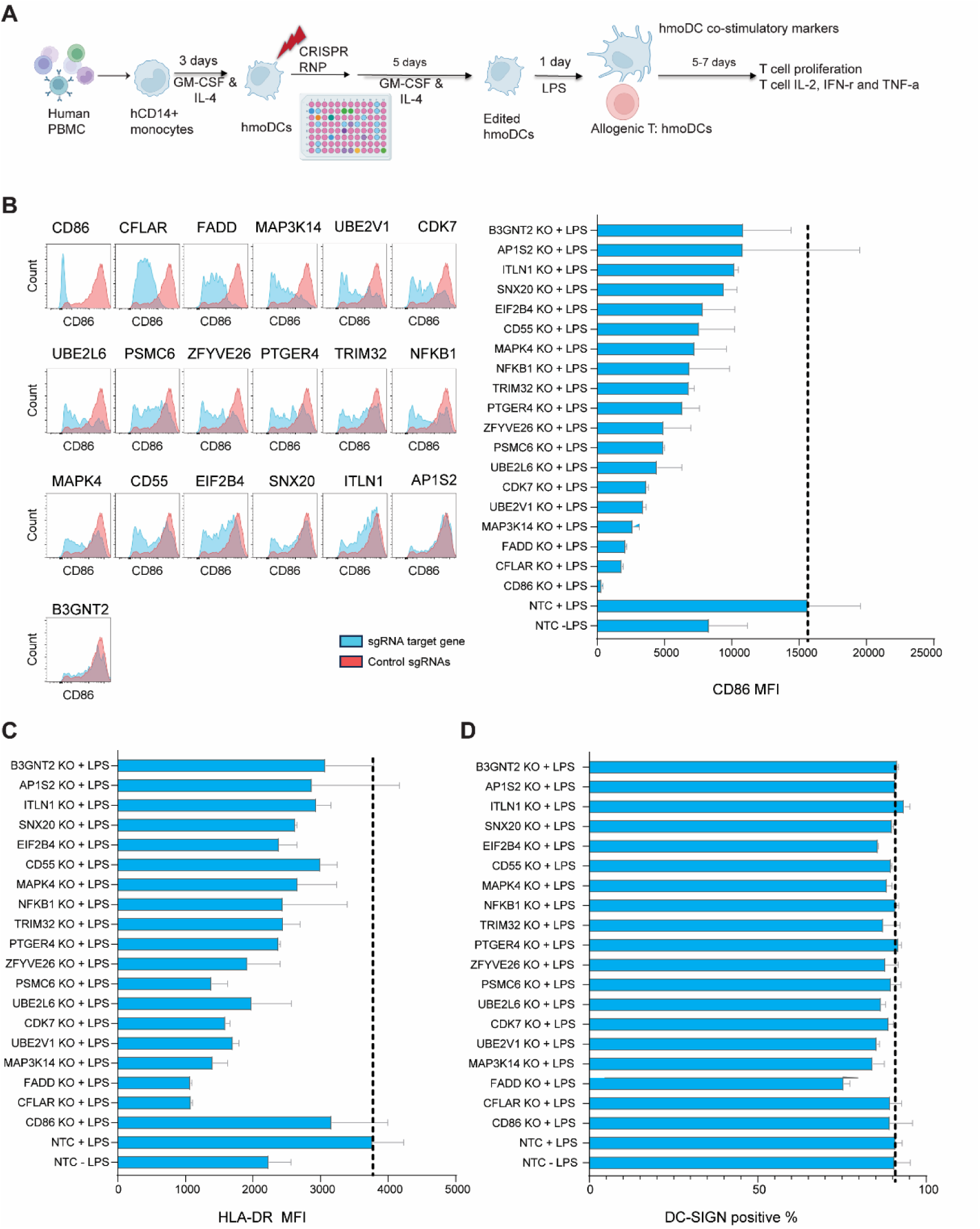
Validation of the screening hits in secondary human CRISPR screen. **(A)** Schematic diagram of CRISPR-Cas9 array screen in human moDCs (hmoDCs). **(B)** Flow cytometry analysis of CD86 expression in moDCs following CRISPR knockout of genes (blue) or non-targeting controls (NTC, red). **(C)** Flow cytometry analysis of HLA-DR expression in moDCs following CRISPR knockout of genes. **(D)** Flow cytometer analysis of DC-SIGN expression in moDCs after CRISPR RNP-mediated KO of selected genes. Representative data from one donor, with N = 3; error bars indicate mean ± SEM.

### UBE2L6 is identified as a critical regulator of DC-mediated T cell activation

The screen uncovered a substantial number of previously uncharacterized TolDC regulators, suggesting novel mechanisms controlling DC mediated tolerance, including several genes involved in protein degradation and ubiquitin pathway. Among these, UBE2L6 (also known as UbcH8), is an interferon-stimulated, bifunctional conjugating enzyme essential for both ISGylation and ubiquitination pathways. To investigate the mechanism by which UBE2L6 deletion leads to reduced CD86 expression, we performed quantitative mass spectrometry analysis in dendritic cells under basal and LPS-stimulated conditions in non-targeting control (NTC) and UBE2L6 KO moDCs (Figure 6A-B). In control cells, LPS stimulation led to increased proteins abundance associated with inflammatory signaling and dendritic cell maturation, such as IL1B, IL6, TNF, PTGS2, NFKB1, NFKB2, and CXCL10. These molecules are key components of the TLR signaling, MAPK pathway, JAK-STAT pathway, Interferon signaling, chemokine signaling pathway, and NF-κB signaling pathway. UBE2L6 expression is significantly elevated under LPS stimulation, suggesting its potential involvement in inflammation (Figure 6B-D). Global analysis showed that most of the ubiquitin–proteasome pathway components, including E3 ligases, DUBs, ubiquitin adaptors, and proteasome subunits, transcriptional factors remained largely unchanged at the protein level upon UBE2L6 loss. UBE2L6 KO resulted in a specific upregulation of ISG15 (a ubiquitin-like modifier involved in ISGylation) and its deconjugating enzyme deubiquitinase (DUBs) USP18, a known negative regulator of type I interferon signaling. Given the previously reported role of USP18 as a negative regulator of innate immune signaling, we next examined downstream signaling pathways relevant to dendritic cell activation. Consistent with attenuated NF-κB activity, modest alterations in IκB protein levels were observed in UBE2L6 -deficient cells. In addition, several transcriptional regulators and signaling molecules were reduced in UBE2L6-deficient cells, including the cytokine subunit IL12B, the transcription factor KLF4, the TNF receptor superfamily member RELT, and the transcriptional coactivator CREBBP (Figure 6E-F). Functionally, UBE2L6 deficient DCs exhibited a reduced ability to promote T cell responses. Co-culture of UBE2L6 KO DCs with T cells led to significantly reduced T cell proliferation and decreased inflammatory cytokines production, including IFN-γ and TNF-a, compared with NTC moDCs (Figure 6G-H).

**Figure 6.**
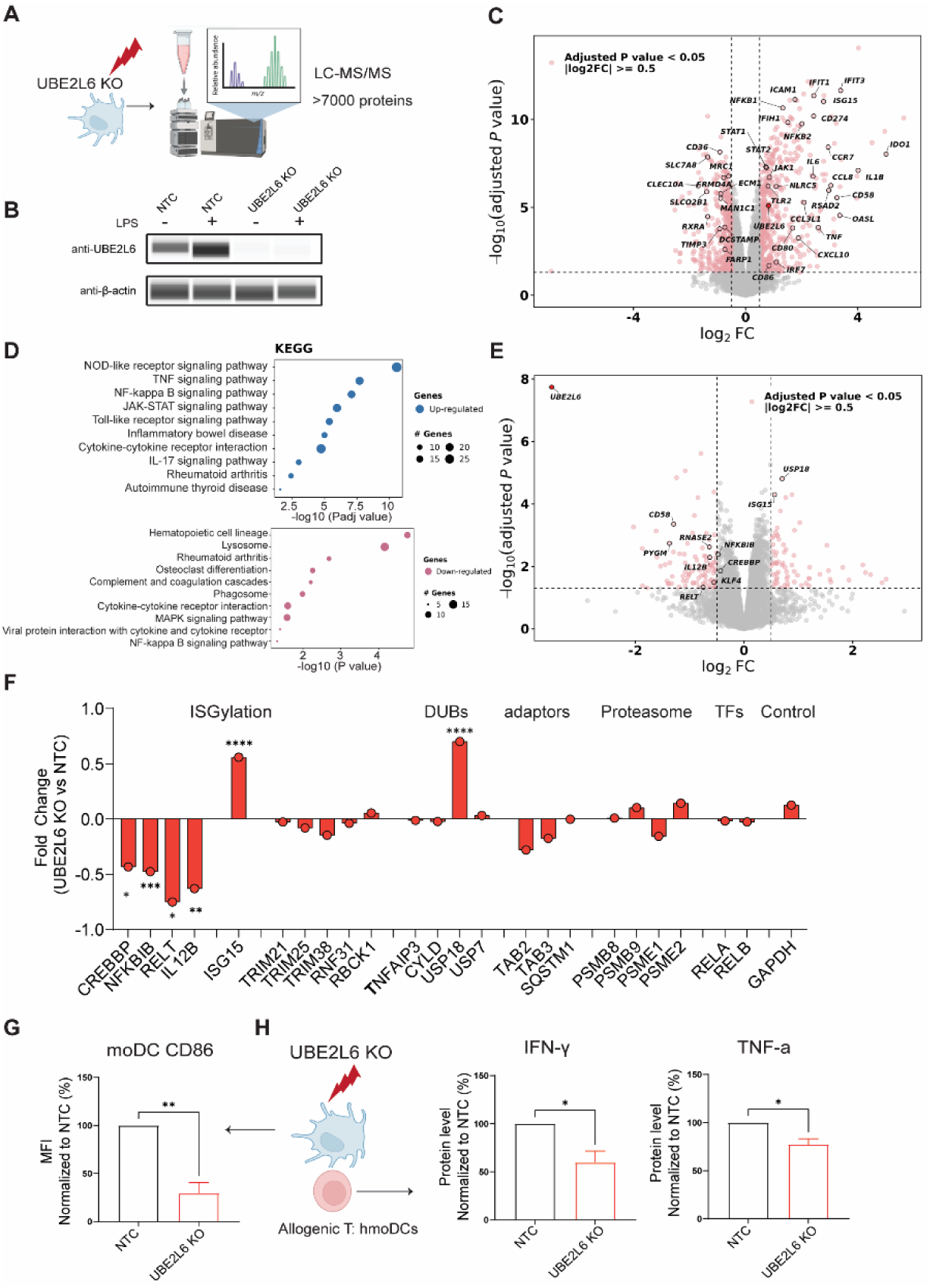
UBE2L6 regulates dendritic cell-mediated tolerance. **(A)** Schematic diagram of LC-MS/MS quantitative proteomics workflow in hmoDCs. (**B)** Representative WES analysis of UBE2L6 expression in UBE2L6 KO and NTC hmoDCs, treated with or without LPS; β-actin as the loading control. **(C)** Volcano plots of global proteomic changes in moDCs treated with LPS vs -LPS vehicle control. Each dot represents one protein. The x-axis shows the log2 fold change (log2FC), calculated as the difference between the mean log2-transformed protein intensities in the +LPS and −LPS groups, y axis depicts log10(Padj-value). **(D)** KEGG pathways enrichment analysis of up-regulated and down-regulated proteins identified by MS from C. **(E)** Volcano plots of global proteomic changes in moDCs between UBE2L6 KO and NTC cells. **(F)** Bar graph of fold change of selected proteins in UBE2L6 KO cells compared with NTC moDCs. **(G)** UBE2L6 was KO in moDCs by CRISPR. Flow cytometer analysis of UBE2L6 expression, normalized to the NTC moDCs. **(H)** Secreted IFN-γ and TNF-α protein level in the supernatant of T cells co-cultured with moDCs, normalized to NTC (%). CD86 expression was measured by flow cytometry, and the cytokine levels measured by MSD assay. The results represent N=3 donors. Statistical significance was calculated with paired two-tailed Student’s t test; error bars show mean with SEM; \**p* < 0.05; \*\**p* < 0.01; \*\*\**p* < 0.001; \*\*\*\**p* < 0.0001.

While not located within a disease-associated GWAS locus itself, the *UBE2L6* gene, which we identified as a novel regulator of TolDC, is regulated by the type 1 diabetes-associated single nucleotide polymorphism rs3184504 through a trans-eQTL effect^36^. Together, these results indicate that loss of UBE2L6 may regulate CD86 expression through induction of the ISG15-USP18 ISGylation axis and attenuation of the NF-κB signaling, thereby impairing dendritic cell activation and DC-mediated T cell priming and inflammatory cytokines.

## Discussion

In this study we established a novel approach to enable identification of mechanisms and therapeutic targets that promote immune tolerance by regulating function of tolerogenic dendritic cells. TolDC are characterized by their abilities to modulate immune responses, suppress effector T cell activity, and promote the generation of regulatory T cells. Progress in this field has been limited by phenotypic heterogeneity and the lack of consensus tolDC markers, as well as by the technical difficulty of applying large-scale functional genomics approaches, including CRISPR-based screens, to primary dendritic cells with limited proliferative capacity. To overcome these limitations and enable comprehensive identification of genes that promote TolDCs, we developed a genome-wide CRISPR screening tailored to TolDC functions, and functionally validating our hits in primary tolDCs. These provide new pathways and molecular targets for enhancing promote immune tolerance in immune-mediated inflammatory diseases.

CRISPR screens have long been powerful tools for assessing gene functions and discovery of novel therapeutic targets that govern the immune response^37–39^. Pooled CRISPR screens have enabled unbiased genome-wide functional comparisons in cell lines that have contributed to the discovery of important pathways and biological mechanisms under various perturbations^31,40,41^. Thus far, very few studies have reported genome-wide functional screens in dendritic cells. Previous studies of CRISPR screens used TNF-α production as a readout of dendritic cell inflammation or activation^29,42^. While TNFα is a well-established marker of pro-inflammatory responses, it does not fully reflect the functional state of tolerogenic DCs, nor does it predict the ability of DCs to modulate downstream T cell responses. Other efforts are restricted to low-throughput arrayed screens in co-coculture assays with allogenic T cells^43^. As a result, genetic architecture that specifically governs tolerogenic DC programs has remained largely unexplored. In this study, we developed a genome-wide CRISPR screening tailored to TolDC functions. Previous reports have shown that TolDCs induced by distinct pharmacological or cytokine-based agents exhibit heterogeneous transcriptional programs, yet these diverse programs ultimately converge on immune tolerance. Comparative transcriptomic analyses across different TolDC-inducing conditions such as VitD3, Rapa, and Dex, revealed a narrow spectrum of consensus markers, suggesting shared regulatory nodes despite diverse upstream signals ^26,44^. Our strategy started with the identification of TolDC intrinsic biomarkers linked to their functional state as there are no lineage-specific TolDC markers associated with TolDC function and TolDC phenotypes assessment relies largely on transcriptomic signatures^44^. We assessed several co-stim molecules or potential consensus markers including CD40, CD86, CD80, IDO-1, MHCII or HLA-DR - using flow cytometer in primary human and mouse TolDCs.

While some markers demonstrated downregulation in specific treatment conditions, CD86 was consistently downregulated across all tested tolerizing conditions, establishing it as a robust and versatile biomarker closely associated with TolDC function. CD86 has been found to be the dominant co-stimulatory molecule for Treg homeostasis and survival as well as regulation of CD4^+^ T cell responses^45^, screens using this biomarker suggest that it is a reliable method to evaluate TolDC function in the clinic. Consistent with these findings, by inactivating CD86 with antibodies or CRISRP/Cas9 editing, we confirmed dependency of a tolerogenic phenotype on CD86. Although CD86 is recognized as interferon and toll receptor stimulated gene, our observation that Dex, VItD3 and Rapa suppressed LPS induced CD86 expression led us to hypothesize that CD86 marks a broader immunosuppressive program of TolDC. Given that genes regulating TolDCs are largely unknown, we performed an unbiased genome wide CRISPR-Cas9 screen in primary DC to comprehensively delineate this program and identify regulators of TolDC phenotype.

To our knowledge, our work represents the first direct genome-wide CRISPR screen in TolDCs, enabling unbiased identification of genes governing tolerance-associated phenotypes. This approach uncovered both canonical immune signaling pathways, such as NF-κB, and previously unappreciated regulators that selectively control CD86 expression, providing new mechanistic insights into TolDC establishment and maintenance. A significant technical challenge in applying pooled CRISPR screening to primary human dendritic cells lies in their limited proliferative capacity and resistance to genetic manipulation. We addressed these technical barriers by optimizing viral delivery conditions and cell expansion strategies, enabling robust library representation and functional screening in primary mouse BMDCs. To demonstrate translatability of mouse model we confirmed that mouse DC cells respond comparably to VitD3 and similarly rely on CD86 downregulation for their suppressive properties. Importantly, we further demonstrated that a substantial fraction of the murine hits showed conserved effects in human monocyte-derived DCs, selectively reducing CD86 expression without broadly impairing DC maturation markers or differentiation. This cross-species validation underscores the translational significance of our results and highlights the therapeutic potential of these regulators for promoting immune tolerance in autoimmune and inflammatory diseases. To assess the relevance of identified targets to inflammation we performed pathway enrichment analysis and found that many targets were linked to critical inflammatory pathways, such as toll-like receptor, IL-17, RIG-I, MAPK, and NFκB signaling, consistent with CD86’s well-established role in immune regulation and disease. To further clarify the inflammatory context and disease relevance of these targets, we examined their associations with autoimmune diseases using genome-wide association studies (GWAS). Many candidate genes showed significant links to increased autoimmune and inflammatory disease risk, indicating that disruptions in dendritic cell co-stimulatory regulation and tolerance pathways may represent a common mechanism in autoimmune pathogenesis.

Notably, we identified UBE2L6 as a key regulator of dendritic cell activation through its control of ISG15 and USP18 protein levels. Proteomic analysis revealed that loss of UBE2L6 specifically increases USP18, a deubiquitinase and central negative regulator of innate immune signaling downstream of Toll-like and cytokine receptors. Both UBE2L6 and USP18 are interferon-stimulated genes, but they serve opposing roles in ISGylation – a process of posttranslational modification of proteins regulating multiple proinflammatory cascades. UBE2L6 adds ISG15 to target proteins, while USP18 removes it. Prior studies and our data suggest that ISG15 may stabilize USP18^46^. Although the exact mechanism behind USP18 upregulation is unclear, we have observed a significant increase in ISG15 levels in UBE2L6-deficient cells.

Therefore, we hypothesize that UBE2L6 deficiency leads to an accumulation of unconjugated ISG15, which in turn stabilizes USP18. Increased USP18 in UBE2L6-deficient dendritic cells may further explain reduced CD86 expression and impaired T cell activation. Besides changes in USP18 we also observed changes in IκB levels which is also known to contribute to tolerogenic state of dendritic cells. Given the central role of USP18 in dampening upstream activation pathways, alterations in IκB dynamics likely reflect reduced signaling flux rather than direct regulation by UBE2L6. In parallel, UBE2L6 -deficient dendritic cells displayed reduced abundance of transcriptional regulators, including CREBBP and KLF4, necessary for NF-κB–dependent gene expression and functional maturation. Loss of these transcriptional cofactors likely compromises the ability of dendritic cells to sustain activation-associated gene programs, including expression of costimulatory molecules such as CD86 and intracellular pools of IL12B. The impaired T cell proliferation and reduced cytokine production observed with UBE2L6 KO DCs may result from downregulation of CD86, NF-κB signaling components (Nfkbib, Relt) and transcriptional regulators (Crebbp, Klf4, Il12b). This suggests that UBE2L6 maintains DC immunostimulatory capacity by regulating key signaling and transcriptional networks necessary for effective T cell proliferation and inflammatory effector function. Collectively, our findings support a model in which UBE2L6 regulates dendritic cell activation by limiting USP18 protein accumulation. Loss of UBE2L6 leads to USP18 stabilization, secondary attenuation of NF-κB–associated signaling, and reduces expression of the costimulatory molecule CD86, ultimately impairing dendritic cell–mediated T cell activation.

By anchoring TolDC function to the robust biomarker CD86, our approach provides insights into the mechanisms underlying immune tolerance that can be used in subsequent high-throughput screens to discover tolerogenic therapies. In addition, our findings set the stage for future efforts to develop effective and safe methods of delivering effector molecules into patient DCs, avoiding *ex-vivo* cell manipulation, laying the groundwork for targeting immune tolerance as a treatment for autoimmune diseases. We envision that future studies should focus on dissecting the downstream signaling and transcriptional networks of identified regulators and evaluating the in vivo relevance of these targets in autoimmune and transplant tolerance models. Our CRISPR-based screening approach, paired with robust functional assays, provides a comprehensive understanding of TolDC biology and powerful toolset for the clinical evaluation and advancement of TolDC-based therapies.

## METHODS

### 1. Differentiation of human monocyte derived DCs and in vitro tolDC generation

Human peripheral blood mononuclear cells were isolated from Leukopaks obtained from Research Blood Components, LLC (USA) using Ficoll gradient-based separation. CD14+ monocytes were magnetically isolated from PBMCs using the StraightFrom® Whole Blood CD14 MicroBeads kit (Catalog# 130-090-879, Miltenyi Biotec, USA). Monocytes were seeded in 6 well plates and cultured in complete RPMI medium at 37 °C in 5% CO_2_. The culture medium was supplemented with 100 ng/ml recombinant human GM-CSFs (Catalog# 300-03-250UG, PeproTech) and 10ng/ml recombinant human IL-4 (Catalog# 200-04-50UG, PeproTech) to differentiate monocytes into moDCs. After 2 days, cells were counted and re-seeded at 50,000 cells/well in 96 well round-bottom plates in complete RPMI supplemented with GM-CSF and IL-4, and treated with concentrations of rapamycin, dexamethasone, or vitamin D3 ranging from 100 nM to 100 pM for 3 days to generate tolerogenic DCs. DMSO was used as a vehicle control. On day 5, cells were washed with 1x D-PBS before stimulating with 100ng/ml lipopolysaccharide (Catalog# tlrl-3pelps, InvivoGen) for 24 hours. Some of the cells were blocked and stained following protocols described in the Flow cytometry Method Section for analysis of co-stimulatory markers.

### 2. Differentiation of bone marrow-derived dendritic cells (BMDCs) and in vitro tolDC generation

Mouse bone marrow cells were isolated from constitutive Cas9 expressing mice (Catalog# 026556, The Jackson Laboratory) or C57BL/6 mice (The Jackson Laboratory). Femur/tibias from mice were harvested and bone marrow flushed out with cold PBS. Cells were filtered through a 70 uM strainer and cultured in complete RPMI medium consisting of RPMI 1640 medium (ATCC Modification) (Catalog# A10491-01, Gibco, USA), 1% Penicillin Streptomycin (Catalog# 15140-122, Gibco), 2mM GlutaMAX (Catalog# 35050-061, Gibco), 1mM Sodium Pyruvate (Catalog# 11306-070, Gibco), 0.1 mM MEM Non-Essential Amino Acids (Catalog# 11140-050, Gibco, USA), 25 mM HEPES (Catalog# 15630-080, Gibco), 55 mM β-mercaptoethanol (Catalog# 21985-023, Gibco) and 10% heat-inactivated FBS (Catalog# 10082-147, Gibco) at 37 °C in 5% CO_2_. Medium was supplemented with 20 ng/mL GM-CSF (Catalog# 315-03-100UG, PeproTech, USA) and 10 ng/mL IL-4 (Catalog# 214-14-100UG, PeproTech) to differentiate cells into BMDCs. After 3 days, cells were treated with concentrations of tolerizing agent ranging from 100 nM to 10^-6^ nM for 5 days to generate TolDCs. On day 8, cells were washed with 1x D-PBS (Catalog# 14190144, Gibco) before stimulating with 100ng/ml lipopolysaccharide for 24 hours. Some of the cells were blocked and stained following protocols described in the Flow cytometry Method Section for analysis of co-stimulatory markers.

### 3. Mixed lymphocyte reaction (MLR) assay

Frozen human PBMCs were thawed and stained with Pan T Cell Isolation Kit, human (Catalog# 130-096-535, Miltenyi Biotec). Stained PBMCs were sorted using autoMACS® Pro Separator (Catalog# 130-092-545, Miltenyi Biotec) according to the manufacturer’s protocols and pan T cells were collected as the negative fraction. Isolated pan T cells were labelled with CellTrace™ CFSE Cell Proliferation Kit (Catalog# C34554, Invitrogen). Mature moDCs or TolDCs and CFSE-stained allogenic pan T cells were co-cultured for 5-7 days in complete RPMI medium at 37 °C in 5% CO_2_. After 5 days, supernatant was collected and secreted cytokine levels were analyzed through MSD assay. The cells were blocked and stained following protocols described in the Flow cytometry Method Section for analysis of T cell proliferation and activation.

OTII CD4+ T cells were isolated from the spleen of OT-II mice using EasySep™ Mouse CD4+ T-Cell Isolation Kit (Catalog # 19852 STEMCELL Technologies) according to the manufacturer’s protocols. Isolated OTII T cells were labelled with CellTrace™ CFSE Cell Proliferation Kit. OVA Peptide (323-339) pulsed mature mouse BMDCs or tolerogenic DCs and CFSE-stained OTII T cells were co-cultured for 5-7 days in complete RPMI medium at 37 °C in 5% CO_2_. After 5 days, supernatant was collected and secreted cytokine levels were analyzed through MSD assay. The cells were blocked and stained following protocols described in the Flow Cytometry Method Section for analysis of T cell proliferation and Treg induction.

### 4. Flow cytometry

Cells were first blocked with either Mouse BD Fc Block™ (Catalog# 553142, BD Biosciences, USA) or Human TruStain FcX™ (Fc Receptor Blocking Solution) (Catalog# 422302, BioLegend, USA) using 1:100 dilution and then stained for 30 min at 4 °C with staining antibodies with 1:200 dilution. DAPI (Catalog# D1306, Invitrogen, USA) or LIVE/DEAD Fixable Aqua Dead Cell Stain Kit (Catalog# L34957, Invitrogen) was used for detecting dead cells. Mouse antibodies for BMDCs include FITC anti-mouse F4/80 Antibody (Catalog# 123108, Clone BM8, BioLegend, USA), PE anti-mouse CD11c Recombinant Antibody (Catalog# 117308, Clone N418, BioLegend), PerCP/Cyanine5.5 anti-mouse CD86 Antibody (Catalog# 105028, Clone GL-1, BioLegend), PE/Cyanine7 anti-mouse CD40 Antibody (Catalog# 124622, Clone 3/23, BioLegend), APC anti-mouse CD274 (B7-H1, PD-L1) Antibody (Catalog# 124312, Clone 10F.9G2, BioLegend), APC/Fire™ 750 anti-mouse I-A/I-E Antibody (Catalog# 107652, Clone M5/114.15.2, BioLegend), Brilliant Violet 510™ anti-mouse CD80 Antibody (Catalog# 104741, Clone 16-10A1, BioLegend), and Brilliant Violet 711™ anti-mouse/human CD11b Antibody (Catalog# 101242, Clone M1/70, BioLegend). Human antibodies for moDCs include FITC anti-human CD83 Antibody (Catalog# 305306, Clone HB15e, BioLegend, USA), PE anti-human CD80 Recombinant Antibody (Catalog# 370612, Clone QA18A16, BioLegend), PE/Cyanine5 anti-human CD86 Antibody (Catalog# 305408, Clone IT2.2, BioLegend), PE/Cyanine7 anti-human HLA-DR Antibody (Catalog# 307616, Clone L243, BioLegend), APC anti-human CD274 (B7-H1, PD-L1) Antibody (Catalog# 329708, Clone 29E.2A3, BioLegend), APC/Fire™ 750 anti-human CD209 (DC-SIGN) Antibody (Catalog# 330116, Clone 9E9A8, BioLegend), and Brilliant Violet 510™ anti-human CD40 Antibody (Catalog# 334330, Clone 5C3, BioLegend). Human T cells were stained with Alexa Fluor® 594 anti-human CD4 Antibody (Catalog# 300544, Clone RPA-T4, BioLegend), Brilliant Violet 421™ anti-human CD8 Antibody (Catalog# 344748, Clone SK1, BioLegend), PerCP/Cyanine5.5 anti-human CD127 (IL-7Rα) Antibody (Catalog# 351322, Clone A019D5, BioLegend), PE/Cyanine7 anti-human CD25 Antibody (Catalog# 302612, Clone BC96, BioLegend), and APC/Fire™ 750 anti-human CD209 (DC-SIGN) Antibody (Catalog# 330116, Clone 9E9A8, BioLegend) using 1:200 dilution. eBioscience™ Foxp3 / Transcription Factor Staining Buffer Set (Catalog# 00-5523-00, Invitrogen, USA) was used according to the manufacturer’s instructions for intracellular staining. Specifically, cells were stained with Alexa Fluor® 647 anti-human FOXP3 Antibody (Catalog# 320114, Clone 206D, BioLegend) and Brilliant Violet 711™ anti-human IFN-γ Antibody (Catalog# 502540, Clone 4S.B3, BioLegend) with 1:50 dilution after fixation.

Samples were recorded on the BD FACSCanto II or BD LSRFortessa flow cytometer (BD Biosciences). Following gating of single cells and exclusion of dead cells, cells were gated as follows: live single cells were gated for DC-SIGN negative cells, which were further gated for FITC negative cells (Proliferated T cells) and APC+/PE-Cy7+ cells (Tregs). FlowJo was used to analyze the data.

### 5. Lentivirus packaging of library

CRISPR Mouse Genome 86K Knockout Library (Catalog# KOMGW-86K-P, Cellecta, USA) were produced by transient co-transfection of the lentiviral vector plasmid and the lentiviral Packaging Plasmid Mix (lentiviral packaging plasmid psPAX2 and the pMD2.G plasmid containing VSV-G) (Catalog# CPCP-K2A, Cellecta) to HEK293T cells (Catalog# CRL-3216, ATCC, USA) using FuGENE® Transfection Reagent (Catalog# E2311, Promega, USA).

Viral supernatants were filtered through 0.45 μm low-protein-binding filters, concentrated by sucrose gradient ultracentrifugation at 107,000g (RCF) for 2 hours at 4 °C (Beckman Coulter Optima XPN-80 using a SW40 Ti rotor). The concentrated lentivirus was resuspended in PBS + 1% BSA, and stored at −80°C.

### 6. Genome-wide CRISPR screening and RNA-sequencing

120 million BMDCs were transduced with the sgRNA library at multiplicity of infection (MOI) = 0.1 in biological duplicates. At 72 h post-transduction, cells were selected on puromycin. Following puromycin selection, cells were treated with LPS for 24 hours. 40 million cells were collected with trypsin as “pre-sort” for genomic DNA extraction for each replicate.

Greater than 120 million cells per replicate were then processed and stained for CD86 at 10 million cells/mL using protocol mentioned in Flow Cytometry section. Cell populations expressing the top and bottom 20% CD86 levels were enriched using a FACSAria Fusion cell sorter (BD Biosciences). Cells were collected for genomic DNA extraction using Quick-DNA FFPE Kit (Catalog# D3067, Zymo Research, USA). PCR reactions containing up to 10 μg genomic DNA in each reaction were performed using ExTaq DNA Polymerase (Catalog# RR001C, Takara Bio, Japan) with primers to amplify the guide RNAs. Samples were then purified with SPRIselect beads (Catalog# B23319, Beckman Coulter, USA), mixed and sequenced on a NextSeq 500 by 75-bp single-end sequencing.

### 7. Sequencing analysis

Sequencing data were demultiplexed and reads containing guide RNA (sgRNA) sequences were quantified using custom Perl scripts. Since the sgRNA could appear at varying positions within reads, the flanking primer sequences were used to locate it. The region between the primers was extracted from each read and matched against the sgRNA library. Only sequences with exact matches (no mismatches) were included when calculating guide-level read counts. Samples were selected for further analysis if at least 80% of their reads were mapped to a guide RNA. Read counts were normalized across samples using the trimmed mean of M values (TMM) method^47^ implemented in the edgeR package^48^.

Differential analysis at the guide level was then performed using the limma package^49^. For gene-level analysis, the median log fold change of the four corresponding guides was used. Statistically significant genes were identified by aggregating differentially ranked sgRNAs at the gene level through the robust rank aggregation (RRA) method^50^. Pathway enrichment analysis was performed using clusterProfiler package^51^.

### 8. Open Targets L2G scores

The Open Targets project has performed systematic fine-mapping and gene prioritization at over 130K published GWAS loci from the GWAS catalog, the FinnGen study and the UK Biobank^52^. The L2G algorithm trained on a set of unequivocally mapped GWAS genes using a gradient boosting model has enabled the ranking of genes within each GWAS locus by integrating statistical fine-mapping and various epigenetic signatures as predictive features. Multiple tables that contain these data from the October 2022 (22.10) release were downloaded in bulk from Google BigQuery^53^. We wrangled the data by merging various tables and harmonizing the phenotype descriptions across different GWAS. We mapped mouse genes to human homologues and then queried the L2G analysis output for the top genes identified in our genome-wide screen CRIPSR. Where more than one study linked genes to the GWAS signals, we selected the highest L2G score among those studies. Top L2G scores between genes and phenotypes above 0.1 were plotted using the *ggplot2* R package (v3.3.6).

### 9. Nucleofection of Cas9/gRNA RNP complex

sgRNAs purchased from Synthego (USA) and Alt-R S.p. HiFi Cas9 Nuclease V3 from (Catalog# 10007803, IDT, USA) were assembled and electroporated into the human moDCs using Lonza P3 Primary Cell 96-well Nucleofector Kit (Catalog# V4SP-3096, Lonza, USA) and optimized program CM-137 in an Amaxa 96-well Shuttle system (Catalog# AAF-1002B, Lonza). Each nucleofection reaction consisted of 0.6-0.8× 10^6^ cells mixed with RNP and 10 uM Alt-R Cas9 Electroporation Enhancer (Catalog# 1075916, IDT). Complete RPMI medium supplemented with GM-CSF and IL-4 for moDC differentiation was added to the cells following electroporation to transfer the cells to 96-well tissue culture plates containing pre-warmed media. MoDCs were cultured for another 4-6 days before treating LPS for DC maturation. The sgRNAs and primers sequences are included in Supplementary Table 1.

### 10. Genomic DNA extraction, PCR, and sanger sequencing

QuickExtract™ DNA Extraction Solution (Catalog# SS000035-D2, Biosearch Technologies, USA) was used to extract gemonic DNA. PCR amplification was performed using Platinum PCR SuperMix High Fidelity (Catalog# 12532-016, Thermo Fisher Scientific) and 10 µM primers (Azenta, USA) for DNA sequencing under the conditions recommended by the manufacturer. C1000 Touch Thermal Cycler (Catalog# 1851196, Bio-Rad, USA) was used to carry out the PCR reaction. PCR amplicons were sent to Azenta for purification and sequencing. Primer sequences are listed in SI Appendix table 1.

### 11. Cytokine detection

Supernatant from cells cultured in 96-well plates was collected, and cytokine concentration was measured with the U-PLEX TH1/TH2 Combo (human) kit (IFN-γ, IL-1β, IL-2, IL-4, IL-5, IL-8, IL-10, IL-12p70, IL-13, and TNF-α) (Catalog# K15010K-4, Meso Scale Discovery, USA). Meso Sector S 600MM instrument (Catalog# IC1AA-0, Meso Scale Discovery) was used to perform the MSD assay and XLfit software (IDBS, UK) was used to analyze the results.

### 12. Protein extraction, BCA assay, and JESS automated Western Blot

Proteins were extracted from cell pellets using RIPA Lysis and Extraction Buffer (Catalog# 89900, Thermo Scientific). Protein concentrations in the lysates were measured by Pierce™ Rapid Gold BCA Protein Assay Kit (Catalog# A53225, Thermo Scientific) and adjusted through comparison to the BSA Protein Assay Standards contained in the kit. 4 parts of lysates were mixed with 1 part of 5X Fluorescent Master Mix from the 12-230 kDa Separation Module (Catalog# SM-W004, Bio-techne, USA) and denatured for 5 minutes at 95 °C. Prepared samples, diluted anti-UBE2L6 primary antibody (Catalog# ab109086, Abcam, USA), anti-beta-Actin primary antibody (Catalog# NB600-532, Novus Biologicals, USA), anti-rabbit secondary antibody, which is provided in the Anti-Rabbit Detection Module (Catalog# DM-001, Bio-techne), and other necessary western blot reagents were placed in the special JESS plate according to manufacturer’s instructions. Automated western blots were performed in the JESS machine (Bio-techne) and results were analyzed in the Compass for Simple Western software.

### 13. LC-MS/MS Analysis and data interpretation

The samples were analyzed using a ThermoFisher Easy nLC 1200 system coupled to an Orbitrap Exploris 480 mass spectrometer. Peptides were eluted with a linear gradient from 6% ACN to 35% Acetonitrile (ACN) over 106 minutes, followed by a 14-minute phase at 90% ACN to wash the column. The Lumos mass spectrometer was operated in DDA mode, with an MS1 scan range from 300–800 m/z at 120,000 resolving power. Fragmentation targeted peptides with charge states between 2 and 7, applying a dynamic exclusion of 25 seconds. Peptides were selected with a 0.8 m/z isolation window and an HCD normalized collision energy of 34%. Tandem mass spectra were analyzed in the Orbitrap at 50,000 resolving power, with an AGC target of 100,000 and a maximum injection time of 120 ms. Data analysis was peformed using Proteome Discoverer (version 2.2, ThermoFisher).

### 14. Statistical analysis

Statistical analysis was performed in GraphPad Prism. one-way ANOVA or unpaired two-tailed Student’s *t*-test were used to calculate statistical significance. The p values were indicated (∗p < 0.05, ∗∗p < 0.01, ∗∗∗p < 0.001, and ∗∗∗∗p < 0.0001).

## Acknowledgements

We would like to acknowledge the AbbVie Genomics Research Center GTECH lab and Vinny Vijaykumar for preparing and sequencing NGS libraries. We would like to thank Arlene Lim from AbbVie Immunology Discovery Research for isolating Bone marrow cells. We want to acknowledge the participants and investigators of the FinnGen study. Figures were generated using Adobe Illustrator and BioRender.

## Author contributions

XL, LC and KK designed and supervised the research; XL, TH, MS, RM, JS, performed research, FR, CG, JW analyzed data, X.L., K. K. wrote the paper.

## List of Supplementary Materials

This PDF file includes:

Figure S1 to S4

Table S1

## Supplementary Materials

**Figure S1:**
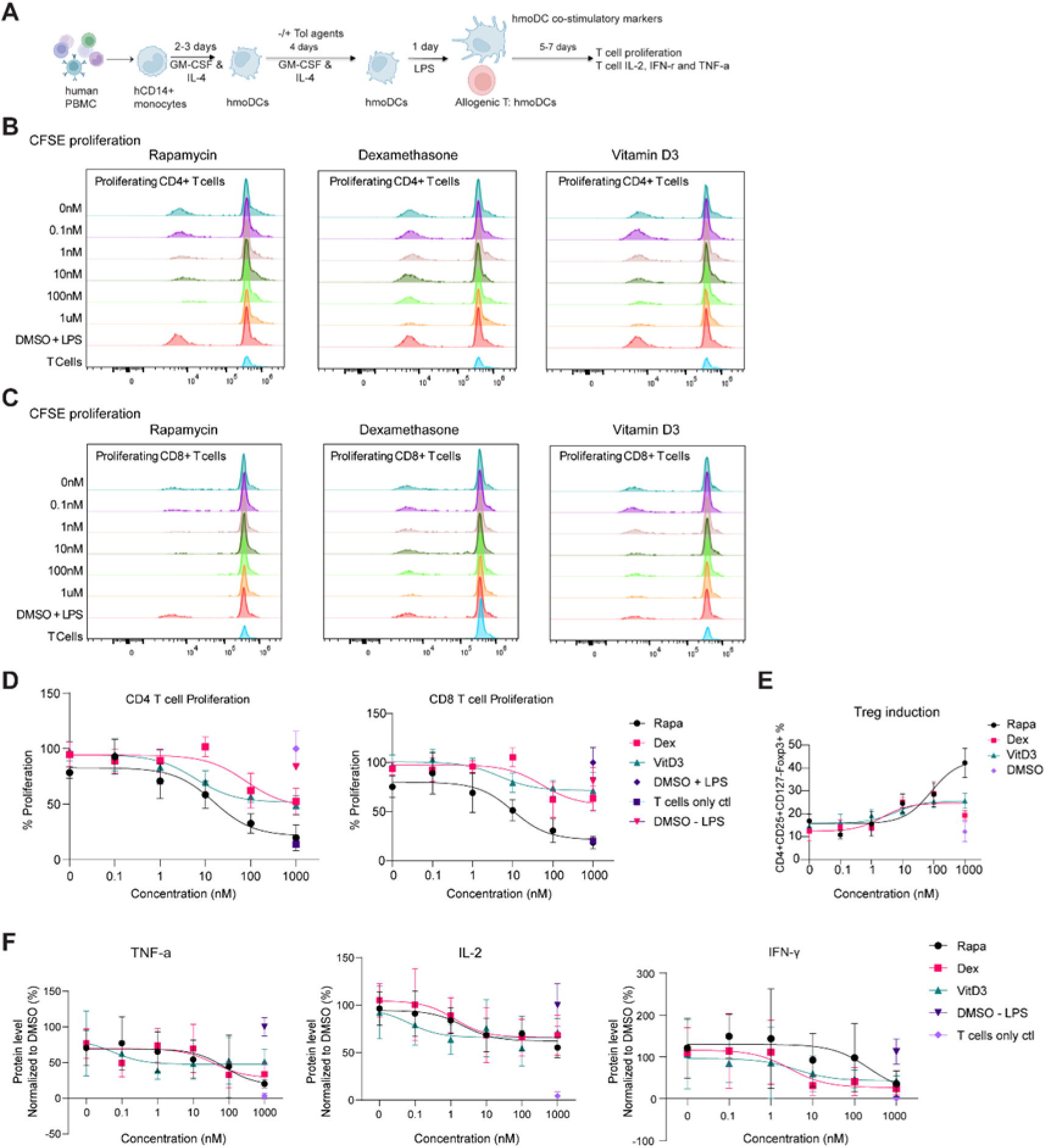
Rapamycin, Dexamethasone, and Vitamin D3 induce tolerogenic phenotypes in human dendritic cells. **(A)** Schematic representation of induction and characterization of human tolerogenic moDCs with treatment of tolerizing agents. **(B–C)** Proliferation of CD4+ T cells (B) and CD8+ T cells (C) co-cultured with tolerogenic moDCs generated using different concentrations of tolerizing agents. Proliferation was assessed by flow cytometric analysis of CFSE dilution. **(D)** Analysis of CD4+ and CD8+ T cell proliferation co-cultured with tolerogenic moDCs as shown in B and C. **(E)** Flow cytometer analysis of frequency of regulatory T cells (Tregs) in the moDC:T cell co-culture, following tolerogenic induction. **(F)** The secreted TNF-α, IL-2, and IFN-γ protein levels in the co-culture with tolerogenic moDCs induced by different concentrations of tolerizing agents. Cytokine levels measured by MSD assay. The results represent N=2 donors in 3 independent experiments. Error bars = mean with SEM.

**Figure S2:**
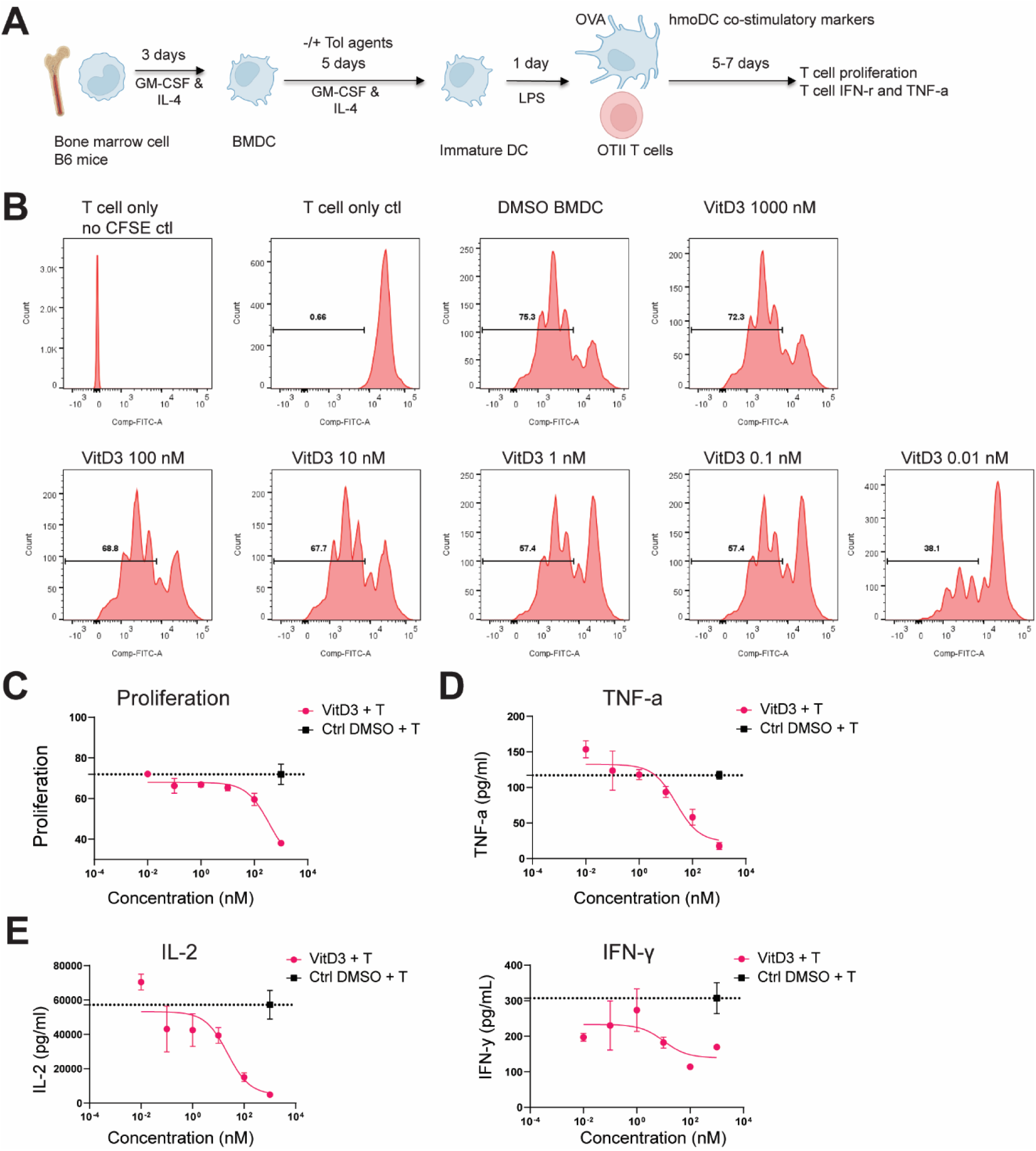
Vitamin D3 induces tolerogenic phenotypes in mouse bone marrow derived dendritic cells. **(A)** Schematic representation of induction and characterization of mouse tolerogenic BMDCs. **(B–C)** Flow cytometer analysis of CFSE dilution demonstrates the proliferation of T cells from OTII mice after co-culture with tolerogenic BMDCs induced by different concentrations of Vitamin D3. BMDC was pulsed with OVA peptide. **(D–E)** The secreted TNF-α, IL-2, and IFN-γ levels in the co-culture assay. Cytokine levels measured by MSD assay. N=3 independent experiments. Error bars indicate mean with SEM.

**Figure S3:**
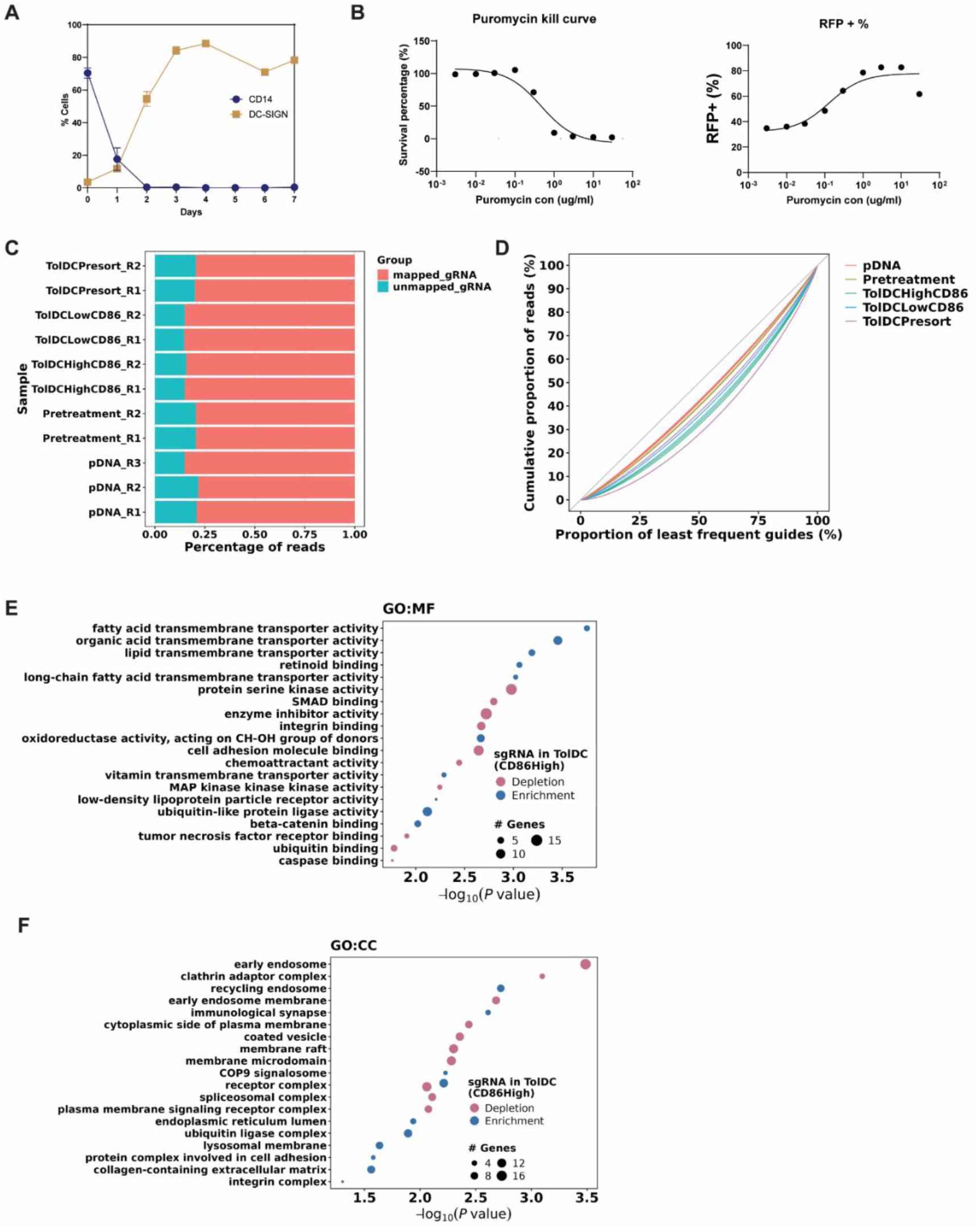
**(A)** CD14 and DC-SIGN expression dynamics during differentiation. Human monocytes/moDCs cultured in medium supplemented with GM-CSF and IL-4. **(B)** Percentage of live mouse BMDCs and RFP-expressing cells in mouse following lentiviral transduction and subsequent puromycin selection at various concentrations. **(C)** Mapping rate of sequencing reads for samples. **(D)** Lorenz curve analysis of sgRNA representation in samples. **(E–F)** Selected terms enriched in positive (red) and negative (blue) regulators of CD86 ranked by - log2(P value) from pathway enrichment analysis using Gene Ontology (GO): Molecular Function (MF) and Cellular Component (CC).

**Figure S4:**
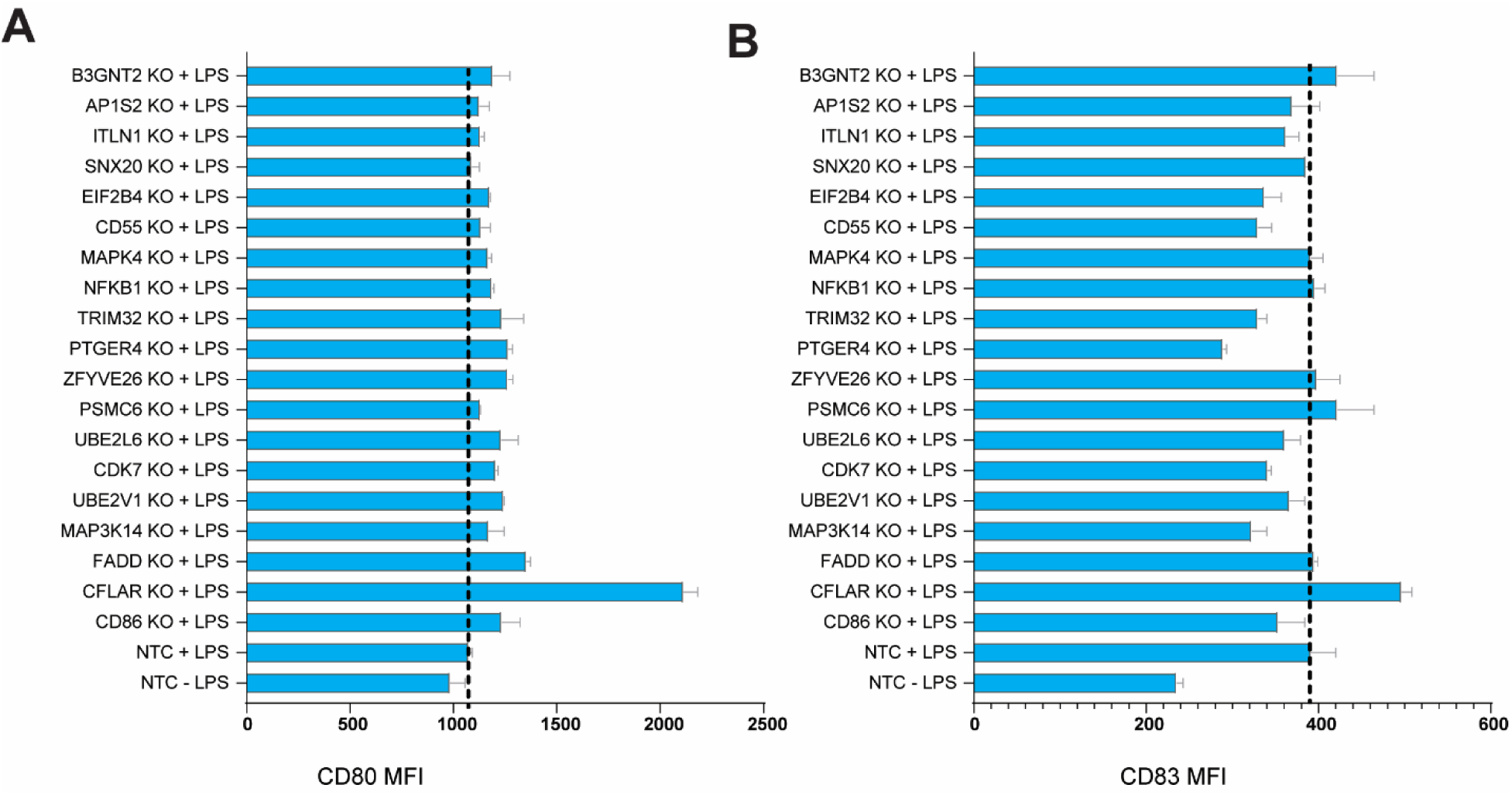
**(A)** Flow cytometer analysis of CD80 expression in moDCs. **(B)** Flow cytometer analysis of CD83 expression in moDCs. The results represent N=3. Error bars indicate mean with SEM.

## Supplementary Tables

**Table S1:**
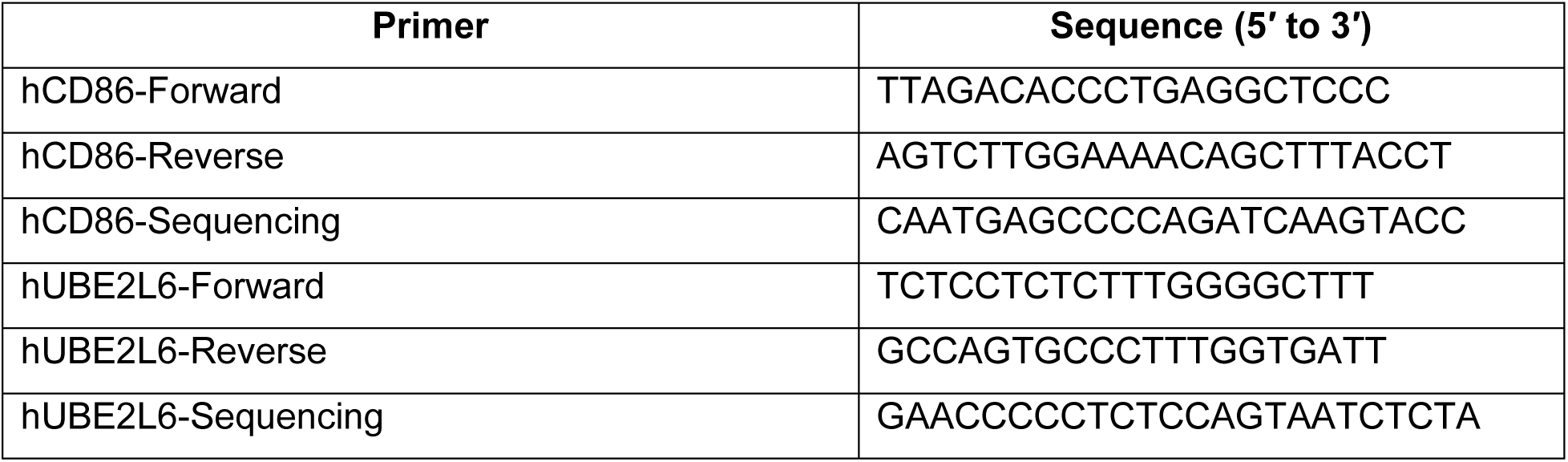
Primer sequences used in the study.

**Table S2:**
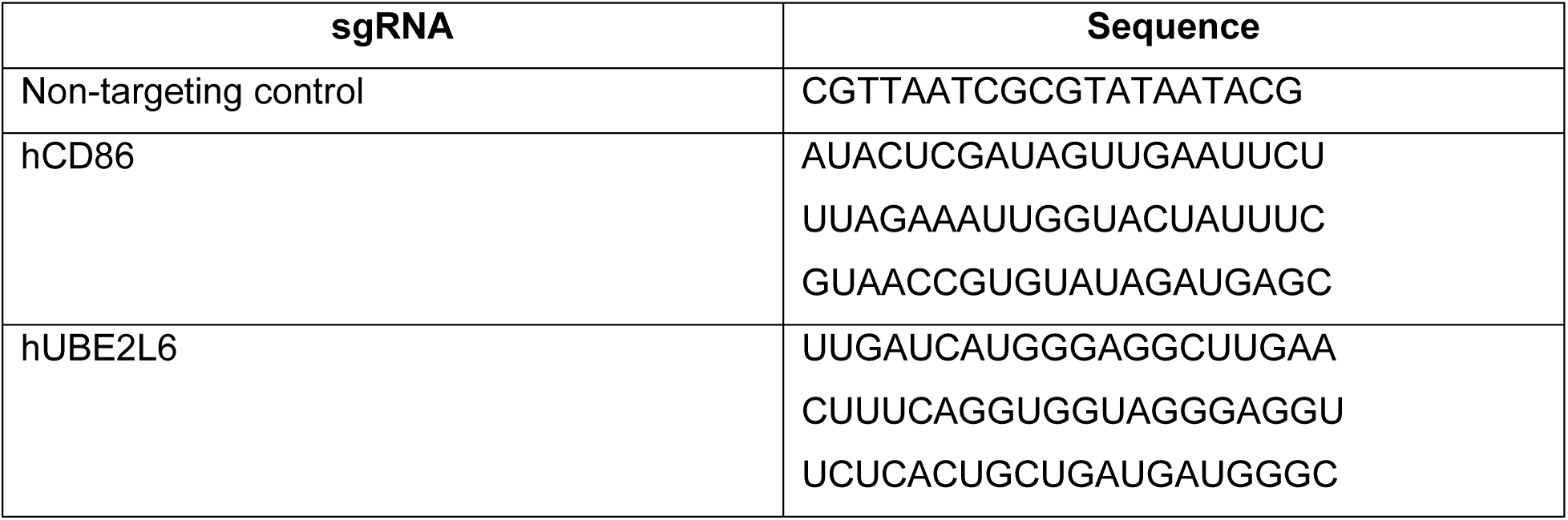
Single guide RNAs (sgRNAs) used in the study.

